# Predicting the Beat Bin: Beta Oscillations Predict the Envelope Sharpness in a Rhythmic Sequence

**DOI:** 10.1101/2023.07.21.550020

**Authors:** Sabine Leske, Tor Endestad, Vegard Volehaugen, Maja D. Foldal, Alejandro O. Blenkmann, Anne-Kristin Solbakk, Anne Danielsen

**Author notes:** Correspondence to: Sabine Leske, RITMO Centre for Interdisciplinary Studies in Rhythm, Time and Motion University of Oslo, Forskningsveien 3A, 0373 Oslo Norway.

## Abstract

Periodic sensory inputs entrain oscillatory brain activity, reflecting a neural mechanism that might be fundamental to temporal prediction and perception. Most environmental rhythms, such as music or speech, however, are rather quasi-periodic. Research has shown that neural tracking of speech is driven by modulations of the amplitude envelope, especially via sharp acoustic edges, which serve as prominent temporal landmarks. In the same vein, research on rhythm processing in music supports the notion that perceptual timing precision varies systematically with the sharpness of acoustic onset edges, conceptualized in the beat bin hypothesis. Increased envelope sharpness induces increased precision in localizing a sound in time. Despite this tight relationship between envelope shape and temporal processing, it is currently unknown how the brain uses predictive information about envelope features to optimize temporal perception.

With the current study, we show that the predicted sharpness of the amplitude envelope is encoded by pre-target neural activity in the beta band (15–25 Hz), and has an impact on the temporal perception of sounds. Using probabilistic sound cues in an EEG experiment, we informed participants about the sharpness of the amplitude envelope of an upcoming target sound embedded in a quasi-isochronous beat. The predictive information about the envelope shape modulated the performance in the timing judgment task and pre-target beta power. Interestingly, these conditional beta-power modulations correlated positively with behavioral performance in the timing-judgment task and with perceptual temporal precision in a click-alignment task.

This study provides new insight into the neural processes underlying prediction of the sharpness of the amplitude envelope during beat perception, which modulate the temporal perception of sounds. This finding could reflect a process that is involved in temporal prediction, exerting top-down control on neural entrainment via the prediction of acoustic edges in the auditory stream.

## Introduction

Humans, among some other species, show a strong tendency to synchronize their movements to periodic events that express a regular pulse or beat (Merchant et al., 2015), and their oscillatory brain activity synchronizes with rhythmic sounds (Obleser and Kayser, 2019). This has been referred to as neural entrainment, and there is an active debate as to whether this entrainment is instrumental to temporal prediction and under active top-down control (Doelling et al., 2023, 2023; Lakatos et al., 2019; Rimmele et al., 2018).

Most environmental rhythms and patterns in human behavior, such as walking, dancing, and speech, however, do not display strict isochrony (Bonnet et al., 2024; Doelling et al., 2023) but are instead quasi-periodic. Nevertheless, neural entrainment to the amplitude envelope (changes in the amplitude of a sound over time) of, for example, speech has been shown to support comprehension (Ding et al., 2017; Giraud and Poeppel, 2012; Lakatos et al., 2019; Poeppel and Assaneo, 2020; Zoefel and VanRullen, 2016). This type of entrainment is also referred to as neural tracking or neural entrainment in the broad sense, in case is not known if the generating mechanism is of oscillatory nature (Obleser and Kayser, 2019). Research supports the conclusion that the sharpness of the amplitude envelope of the speech signal works as an acoustic landmark or “edge” and is a critical factor in driving neural tracking, improving both syllabic segmentation and intelligibility (Doelling et al., 2014; Ghitza, 2013). Envelope sharpness has been defined as rapid changes in the amplitude envelope that correspond to acoustic onset edges (Oganian and Chang, 2019) that have a short rise time, also referred to as a sharp attack (Irsik et al., 2021). We will use the term attack, when referring to rise time throughout the paper. Envelope sharpness can be calculated as the positive first derivative of the amplitude envelope (Doelling et al., 2014; Oganian and Chang, 2019). Deficits in the detection of the attack and impaired audio-motor synchronization to a beat are apparent in developmental dyslexia (Colling et al., 2017; Goswami, 2011; Goswami and Leong, 2013). This provides additional evidence that the sharpness of the speech envelope is critical to rhythm synchronization and language comprehension (Colling et al., 2017; Goswami, 2011; Goswami et al., 2002; Huss et al., 2011; Power et al., 2016; Woodruff Carr et al., 2014).

Likewise, research on the perception of timbre and source identity has shown that when attack information is removed from the sound, the perceiver’s ability to identify the instrument decreases drastically (Berger, 1964; McAdams and Siedenburg, 2019; Rasch and Plomp, 1999; Saldanha and Corso, 1964). These results suggest that acoustic edges, and in particular the shape of the attack, carry important information about timbre and source identity in music perception.

Importantly, research into human perception of musical rhythm has demonstrated that the perceptual timing precision of a sound depends on the acoustic envelope shape (Danielsen, 2010; Danielsen et al., 2019). For example, a short sound with a sharp attack is perceived with high temporal precision, whereas a long sound with a gradual (i.e., smooth) attack, like a violin note, gives rise to a perceived pulse that allows for some rhythmic flexibility and features lower temporal precision (Danielsen et al., 2019). Previous research thus points to a tight link between the sharpness of the amplitude envelope and the precision of beat synchronization.

If neural tracking (or entrainment) reflects a predictive mechanism subserving the temporal prediction and perceptual processing of quasi-periodic sounds, then the expected shape of the amplitude envelope plays a crucial role in the top-down control of entrainment. A respective neural process could therefore ensure a dynamic adjustment of neural entrainment to expected deviations from periodicity and important acoustic landmarks, possibly via a phase reset of oscillatory activity. Here, we ask if the predicted sharpness of the amplitude envelope in a rhythmic sequence is used for such dynamic adjustments, affecting temporal perception. The goal of the current study was to reveal neural mechanisms underlying predictions about the envelope shape of periodic sounds, thereby indirectly addressing how the brain prepares for different levels of perceptual temporal precision.

The perceived temporal precision of a sound can be measured via its perceptual center location (P-center) and its variability (Danielsen et al., 2022, 2019; London et al., 2019). The P-center reflects the subjectively perceived time of occurrence of a sound (Morton et al., 1976). This can vary substantially for different sound types, leading, for example, to high precision (for a sharp sound) or low precision (for a smooth sound) in the perception of its time of occurrence across instances (Danielsen et al., 2022, 2019; Gordon, 1987; London et al., 2019; Villing, 2010; Vos and Rasch, 1981). It has been shown that the variability of the P-center decreases with increasing sharpness of the attack of musical tones (Gordon, 1987; Wright, 2008). In music cognition studies, this has been referred to as the beat bin (Danielsen et al., 2022; London et al., 2019), which denotes the concept that the perceived beat is not defined as a definite point in time but rather by a probability distribution. Hence, the beat bin reflects rhythmic tolerance or the temporal window within which sonic events can occur and still be perceived as (part of) the beat (Danielsen et al., 2022, 2019; Johansson, 2016). The temporal precision of beat perception seems to vary systematically with amplitude envelope features such as attack and duration (Danielsen et al., 2019), with increasing sharpness of the attack and decreasing duration inducing increased perceived beat precision. Accordingly, we will use the term perceptual timing precision when referring to the width of the beat bin throughout the rest of this paper.

Systematic investigations of the effect of the amplitude envelope sharpness on neural entrainment are largely missing (but see Doelling et al., 2019, 2014); exceptions include those studies investigating neural tracking of the amplitude envelope during speech perception (Kösem and van Wassenhove, 2017; Oganian and Chang, 2019; Poeppel and Assaneo, 2020) and a study showing that auditory-motor synchronization can be modulated by acoustic features in a subgroup of participants (Mares et al., 2023). But the emphasis of speech tracking studies is often to control for differences in acoustic properties. This is surprising, given that the input shape can have a major impact on neural synchronization because oscillators are not shape invariant (Doelling and Assaneo, 2021). However, the nature of this relation is still unclear.

The phase-reset of ongoing spontaneous slow (< 10 Hz) oscillations is typically considered to be the mechanism underlying neural entrainment to external rhythms (Henry et al., 2014; Henry and Obleser, 2012; Kösem et al., 2014; Lakatos et al., 2019; Large and Jones, 1999). However, neural entrainment might also manifest as the alignment of rhythmically generated cortical oscillatory bursts entailing higher frequency ranges (Lakatos et al., 2019). While neural entrainment can arise directly from environmental inputs, these inputs can also be linked to rhythmic motor sampling patterns that reset and entrain neural activity in sensory areas (Lakatos et al., 2019; Merchant et al., 2015). This neural process has been referred to as active sensing (Lakatos et al., 2019; Rimmele et al., 2018; Schroeder et al., 2010).

In support of the active sensing framework, Morillon et al. (2014, 2017) revealed that rhythmic movement enhances acoustic temporal perception. Beta (∼20 Hz) activity originating in the sensorimotor cortex was time-locked to the expected auditory targets and directed toward auditory brain regions while being coupled to the phase of delta oscillations (1–3 Hz) at the beat rate of the stimuli (Morillon and Baillet, 2017). Moreover, Fujioka et al. (2015, 2012) reported that beta activity in interacting motor and auditory regions in humans predicts the timing of an auditory beat already before its onset. Beta activity might thus represent the internalization of predictable intervals of a beat and reflect the efference copy signal from the motor system generated at the tempo of the beat (Fujioka et al., 2015, 2012; Merchant et al., 2015). If neural beta activity plays a functional role in auditory beat prediction, then it should be modulated in a top-down manner by predictions about the envelope shape.

It has been proposed that one role of beta bursts may be to phase reset slower oscillations, aligning excitability phases with the time of occurrence of expected stimuli and thus optimizing perception at a low metabolic cost (Canavier, 2015; Sadaghiani et al., 2022). However, mechanisms underlying this type of active sensing, and the respective neural pathways controlling the gain of beat perception, are not fully known (Morillon et al., 2019).

In the present study, we examined whether predicting the sharpness of the amplitude envelope affects the temporal processing of sounds. We manipulated envelope sharpness by varying the duration of the attack (positive derivative of amplitude envelope) and the total duration of the sound. In particular, we targeted neural processes supporting the prediction of different levels of sharpness and, in turn, the prediction of different levels of temporal precision in beat perception. Participants heard a sequence of isochronous sounds and had to judge the relative timing of a target sound at the end of the sequence as either “on time” or “delayed”. We built on the findings of human rhythm-perception research reported above by employing acoustic features (attack, duration and center frequency) that create envelope shapes of different degrees of sharpness that have been shown to induce high or low temporal precision in beat perception (Câmara et al., 2020; Danielsen et al., 2022; London et al., 2019, 2019). By manipulating the listener’s expectation for the envelope’s sharpness (sharp sound equals sharp attack and short duration; smooth sound equals gradual attack and long duration) of the target sounds via sound cues, we sought to induce different predictions regarding the perceptual timing precision (high or low) of a target sound following an isochronous beat sequence (Figure 1). The sound cue was valid in the majority of the trials to enable the reliable prediction of the envelope shape of the target. As an additional control, and to rule out the possibility that neural responses are purely bottom-up and stimulus driven, we investigated predictions solely based on inferred cue-to-target transition probabilities from previous trials (i.e., the inferred occurrence probability of a specific target given a particular cue). Neural responses to transition probabilities reflect a top-down contribution to the prediction process. Participants performed a timing-judgment task that in turn indicated whether the cue information was behaviorally relevant and utilized in the temporal prediction processes of the brain.

In accordance with studies investigating temporal prediction and beat perception (Arnal et al., 2015; Foldal et al., 2023; Fujioka et al., 2015, 2012; Kulashekhar et al., 2016; Merchant et al., 2015; Morillon and Baillet, 2017; Rimmele et al., 2018), we hypothesized that cue validity would affect performance in the timing-judgment task (Ede et al., 2012; Golob et al., 2002; Nobre and van Ede, 2023; Posner et al., 1980), especially for sharp target sounds, given that these require high perceptual timing precision. Furthermore, we anticipated enhanced pre-target beta power when participants expected a sharp sound (relative to a smooth sound), supporting predictions of high perceptual timing precision. This might reflect a neural mechanism subserving temporal prediction, controlling neural entrainment in a top-down manner.

## Results

We reasoned that if neural tracking to quasi-isochronous sounds with a varying envelope is under top-down control, then a cue indicating the envelope sharpness of a target sound should affect temporal perception—that is, the judgment of the timing (delayed or on time) of the target sound embedded in a beat. We used a two-alternative forced-choice timing-judgment task to test whether cue validity affects detection performance. Participants (n = 19) had to detect the timing of a target sound and responded via button press (indicating delayed or on time). Importantly, the target sound followed three isochronous entraining sounds that were physically constant across conditions and presented at a rate of 1.25 Hz (see Figure 1).

**Figure 1:**
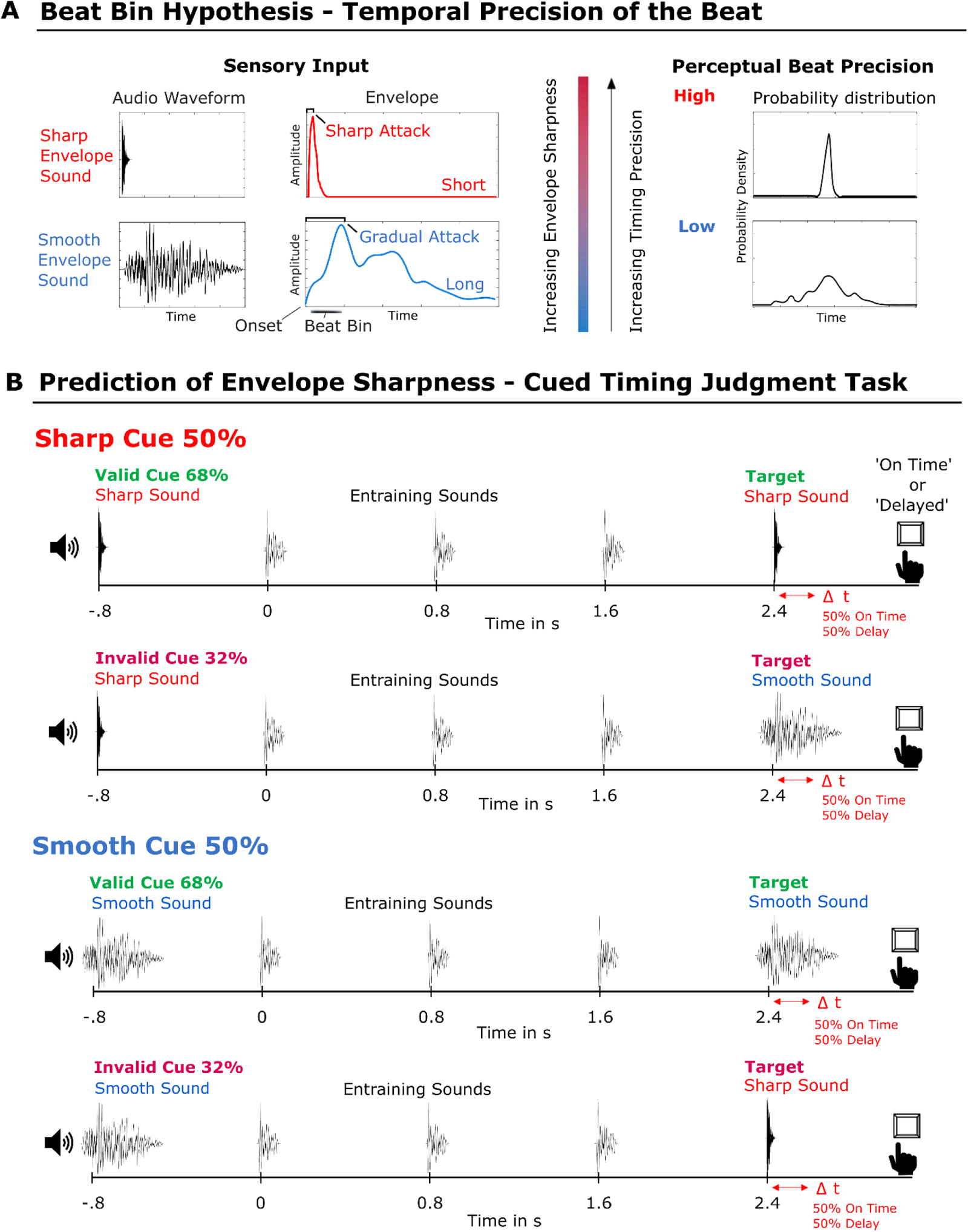
Hypothesis and Experimental Design. (A) Exemplary illustration of the beat bin hypothesis. Schematic illustrations of sensory input (Left) and induced perceptual beat precision (Right). The sharpness of the amplitude envelope shape of the sound modulates the temporal precision of the perceived beat. The acoustic shape of the sharp sound (red, short attack and short duration) induces a high-precision beat percept. The acoustic shape of the smooth sound (blue, gradual attack and long duration) induces a low-precision beat percept, which allows for enhanced flexibility with respect to perceived beat timing. The beat bin reflects the temporal width of the beat percept. Schematic probability density distributions for both sound types (Right) resulting from a synchronization task. The probability density distribution represents the variations across instances while participants localize a rhythmic event in time. A smooth envelope sound induces low perceptual timing precision, reflected in a wide and flat probability density distribution, and a sharp envelope sound induces high perceptual timing precision, reflected in a narrow (small standard deviation) probability density distribution. (B) Experimental design: two-alternative forced-choice timing-judgment task incorporating a sound cue paradigm. Shown are all factor combinations: envelope sharpness, cue validity, and target timing (2 x 2 x 2). Participants were asked to judge whether the target sound was delayed or on time relative to an isochronous sequence of three entraining sounds. Target sounds were on time or delayed with 50/50% probability. The sharp sound cue indicated a sharp target sound, possibly inducing the prediction of high perceptual timing precision (red). The smooth sound cue indicated a smooth target sound, possibly inducing the prediction of low perceptual timing precision (blue). Stimulus plots are a visualization of the waveforms of the sounds used in the study: the sharp sound (red), the smooth sound (blue), and the entraining sound (black). The cue validity was 68%. In the case of an invalid cue (32%), the smooth target sound was presented for the sharp sound cue and vice versa. The slightly earlier onsets of the entraining and smooth sounds schematically illustrate the P-center alignment in the study.

The trials started with a sound cue indicating the identity of the upcoming target sound that varied with respect to three acoustic features (attack, duration and center frequency). By this, we aimed to manipulate expectations regarding the envelope sharpness of the target (sharp or smooth sound) and thereby induce the prediction of high or low perceptual timing precision. The cue preceded three entraining beats by approximately 800 ms and was either a sharp sound indicating the same sharp sound as a target or a smooth sound indicating the respective smooth target sound (68% cue validity) after three entraining sounds. In the case of an invalid cue (32%), the opposite target was presented.

### Performance in the timing judgment task is affected by the predicted envelope sharpness

To ensure an isochronous beat perception of 1.25 Hz between sounds with different degrees of envelope sharpness, the sounds were aligned according to individual P-center locations measured via a click-alignment task at the beginning of the experiment (see “Methods”). First, we tested whether the stimuli showed the expected perceptual effect with respect to perceptual timing precision, which is reflected by the P-center variability measured via the standard deviation (std) across trials (Figure 2). We replicated the directionality of the results of Danielsen et al. (2019) and Câmara et al. (2020) showing that the sharp sound had the earliest P-center location relative to sound onset (group mean: 6.84 ms) and the smallest variability (group std: 29.48 ms), followed by the entraining sound (group mean: 18.28 ms, group std: 42.77 ms), and that the smooth sound had the highest mean values for the P-center location (group mean: 50.99 ms) and variability (group std: 62.41 ms). Signed-rank tests showed that the two sound types (sharp and smooth) differed significantly from one another in their P-center location (Z = 4.04, P = 0.00003, Cohen’s d = 1.6319) and variability (Z = 3.18, P = 0.0007, Cohen’s d = 0.8271, Figure 2). This result is important because it ensured that the acoustic features (attack, duration and center frequency) used for the two cue conditions (sharp versus smooth) had the intended effect on the perceptual timing precision (inverse of variability).

The sharp sound did not differ significantly from the entraining sound in P-center location (Z = 0.93, P = 0.175) but did differ in P-center variability (Z = 1.927, P = 0.027, Cohen’s d = 0.4467). In contrast, the smooth sound differed significantly from the entraining sound in P-center location (Z = 3.24, P = 0.0006, Cohen’s d = 1.2499) but did not differ significantly in P-center variability, showing only a trend effect (Z = 1.61, P = 0.0542). This confirmed that the perceptual effect (P-center location and variability) of the acoustic features of the entraining sound fell between those of the sharp and smooth sounds. The target delay thresholds, estimated from the training block to ensure above-chance-level performance in the timing-judgment task, also differed significantly between the sharp (threshold group mean: 100 ms) and smooth sounds (threshold group mean: 110 ms), with a significantly higher mean for the latter (Z = 3.33, P = 0.0004, Cohen’s d = 1, Figure 2).

We calculated the d-prime measure to estimate performance and response bias (i.e., tendency to respond either on time or delayed) in the timing-judgment task for the factors envelope sharpness (sharp/smooth) and cue validity (valid/invalid). A repeated-measures ANOVA with those factors showed a significant main effect of cue validity (F(1,18) 6.6519, P = 0.0189) but not of envelope sharpness (F(1,18) = 0.7797, P = 0.3889), and it did not show a significant interaction between cue validity and envelope sharpness (F(1,18) = 2.0957, P = 0.1649). Dependent-samples t-test showed that perceptual sensitivity (d-prime) was significantly increased for validly cued sharp targets (valid-sharp cue condition) compared to invalidly cued sharp targets (invalid-smooth cue condition) (t(18) = 2.4593, P = 0.0121, Cohen’s d = 0.5642). This indicates that performance in timing judgment deteriorated when the participants expected a target sound with a low envelope sharpness but received a target sound with a high envelope sharpness (Figure 2). In contrast, there was no effect of cue validity for trials with a smooth target sound, as there was no significant difference between validly cued smooth and invalidly cued smooth targets with respect to the d-prime measures (t(18) = 0.0961, P = 0.4622). Moreover, there were no significant differences in the response bias for any of the conditions (data not shown).

**Figure 2:**
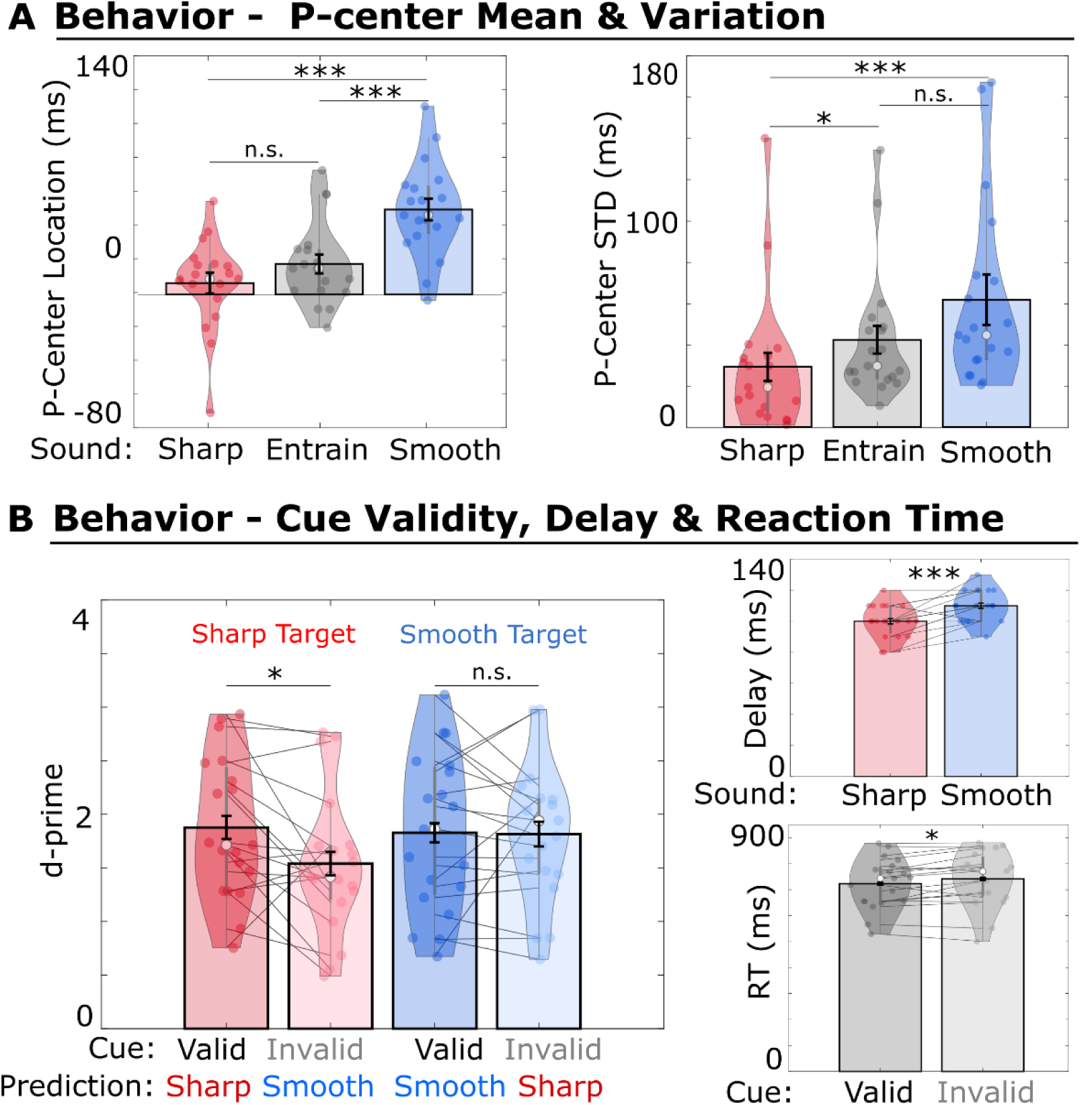
Behavioral Results. Bar plots represent the mean and black error bars represent the adjusted standard error (SE) across participants for a within-subject design, according to the Cousineau-Morey correction (O’Brian & Cousineau (2014)). Violin plots show kernel density estimates and data points (Berchant, 2016). Asterisks refer to significant differences between conditions (*< 0.05, **< 0.01, ***< 0.001). (A) P-center location and variability (std) for all stimuli, sharp (red), entraining (gray), and smooth (blue) envelope sound from the click-alignment task. (B) Behavioral results for the timing judgment task. Left: Valid cue conditions are depicted in dark colors, and Invalid cue conditions are depicted in light colors. Sensitivity (d-prime) was relatively increased for the valid-sharp cue condition (dark red) compared to the invalid-smooth cue condition (light red) (i.e., when participants unexpectedly received a sharp target sound). Sensitivity was not affected by cue validity for the smooth target trials (valid-smooth cue condition [dark blue] versus invalid-sharp cue condition [light blue]). Upper Right: Individual delay threshold results from the training block preceding the main experiment. These thresholds were used for the main experimental blocks. Lower Right: Reaction times for the valid versus invalid cue conditions (pooled over the factor levels envelope sharpness and target timing).

Altogether, these results indicate that the cue information about the envelope sharpness of the forthcoming target sound was behaviorally relevant to temporal perception (Figure 2). An ANOVA showed a significant main effect of cue validity on reaction times (F(1,18) = 5.7866, P = 0.0271, Cohen’s d = -0.5519). Since one of the eight factor levels for the ANOVA deviated from normality, a non-parametric signed-rank test was performed for valid cue trials versus invalid cue trials (pooled over envelope sharpness and target timing, Figure 2), confirming a significant effect of cue validity on reaction time (z-value = -2.1328, p-value = 0.0329). Comparisons did not achieve significance when performed separately for the two envelope-sharpness conditions.

Interestingly, most of the participants (79%) stated in the post-experiment self-report that they did not consciously use the information in the cue to solve the task. Participants were instructed not to move during the experiment; 47% reported that they did not move to the beat at all, 37% reported that they moved to the beat on some occasions, and 16% reported that they moved to the beat during the experiment. The majority (68%) of the participants indicated that they had the impression that it was easier to judge the timing of the sharp target sound. See “Supplementary Information” for the other results of the questionnaire.

### Beta-band power is modulated by the predicted envelope sharpness

To reveal the neurophysiological processes underlying the prediction of envelope sharpness, we focused on pre-target beta oscillations during the entrainment interval, aligned to the first of the three entraining sounds. We assumed that the presentation of sound cues would induce different levels of prediction of the perceptual timing precision of the target sound that appeared at the end of the beat sequence. Accordingly, we concentrated on the main factor of envelope sharpness, comparing trials starting with a sharp sound cue to trials starting with a smooth sound cue.

Grand average spectra across participants were calculated for the entire entrainment period (0–2.4 s). The spectra revealed narrowband peaks of power in the theta (4–6 Hz), alpha (8–12 Hz), and beta (15–25 Hz) frequency ranges, indicating the involvement of oscillatory activity (Supplementary Figure 1).

The temporal modulation of the beta-power time series revealed a rhythmic alignment of beta power to the entraining sounds for the sharp cue condition at certain electrodes. For example, beta-band power at left (C5, T7) and right (C4, C6, T8) centrolateral electrodes evinced an Event-Related Depression (ERD) following the entraining sounds and a beta-power increase (Event-Related Synchronization, ERS) shortly before the next entraining sounds (Figure 3). Also other electrodes, such as frontal (F2, F3, F5, FCz), parietal (P8, P10, CP5, CP6, TP8) and occipital sensors (Oz, Iz) showed a beta-power decrease after and an increase shortly before the entraining sounds, apparently tracking the timing of the sound sequence for the sharp cue condition. Beta-power time series showed different dynamics depending on the cue that were already evident after the first entraining sound and throughout different time segments of the entire entrainment period (Figure 3).

For the following analysis, we concentrated on the time period shortly before target onset, starting with the last (third) entraining sound and ending with the first possible target sound onset (see “Methods” for details). This corresponds to the time period 1.6–2.28 s relative to the first entraining sound. Based on previous studies investigating beat perception and temporal processing (Arnal et al., 2015; Fujioka et al., 2015, 2012; Fujioka and Ross, 2017; Graber and Fujioka, 2020; Morillon and Baillet, 2017), which reported a positive relationship between beta power and perceptual timing performance or predictive timing, we also expected a beta power increase for the sharp cue condition in our data. The statistical contrast (one-sided cluster-based permutation test, n = 19) between sharp versus smooth cue conditions revealed that beta power was significantly enhanced for the cue condition predicting a sharp target sound. Depending on the electrode, this relative increase in beta power started at the earliest approximately 500 ms prior to the target sound and lasted at the most until the earliest possible occurrence of that target sound (range = 1.7750–2.2800 s), entailing frontal, central, temporal, parietal, and occipital sites (cluster T-sum = 3172.5919, cluster P-value = 0.0282). The effect size for the average power values across channels and time points was Cohen’s d = 0.8353.

To control for differences in the physical features between the sharp and smooth cues at the beginning of the trial and a possible confounding effect on pre-target beta power (2.4 s later), we analyzed transitional probabilities to contrast expected sharp versus smooth targets based on the history of cue-target transitions. Sharp and smooth cues can be followed by sharp or smooth targets with different levels of probability (valid and invalid cue-target pairs). Given the random nature of the experiment, fluctuations in the occurrence probability of different cue-target pairs (valid and invalid) are to be expected within short time windows. It has been shown that the brain can track random fluctuations of transitional probabilities (Fuhrer et al., 2023; Maheu et al., 2019; Meyniel et al., 2016). Therefore, we hypothesize that, if there is a top-down mechanism predicting the envelope sharpness for a specific target, then the expected transition to specific targets, based on the short history of recent cue-target pairs, should modulate beta-power oscillations in the same direction as the cue conditions. In other words, if for a short period sharp cues are mostly followed by sharp targets, pre-target beta power should increase in the following trials, but if sharp cues are mostly followed by smooth targets (i.e., invalid sharp cues), pre-target beta power should decrease. In practical terms, we modeled the probability of receiving a sharp target sound for each trial by taking into account valid and invalid cues of previous trials (for details, see “Methods”). This allowed us to contrast the high (75th or upper quartile) versus low (25th or lower quartile) probability trials to receive a sharp target sound for each cue condition (sharp and smooth) individually. Beta-power time series were modulated in the same direction (Figure 3B), with increased pre-target beta power for a high (75–100%) transition probability (TP) to receive a sharp target, as opposed to the low TP (1–25%) to receive a sharp target sound (likely inducing an enhanced expectation to receive a smooth target). This effect was apparent for the sharp cue condition (cluster T-sum = -2651.3616, cluster P-value = 0.0410) between 1.865–2.28 s for parietal sensors. The effect size for the average power across channels and time was Cohen’s d = 1.3827. For the smooth cue condition, the high versus low TP contrast was not significant.

Beta-power time series preceding the target by approximately 800 ms (1.48–2.28 s, to avoid evoked responses by early target onsets in some participants; see “Methods”) were submitted to a spectral decomposition to investigate frequency modulations of beta-power dynamics.

**Figure 3.**
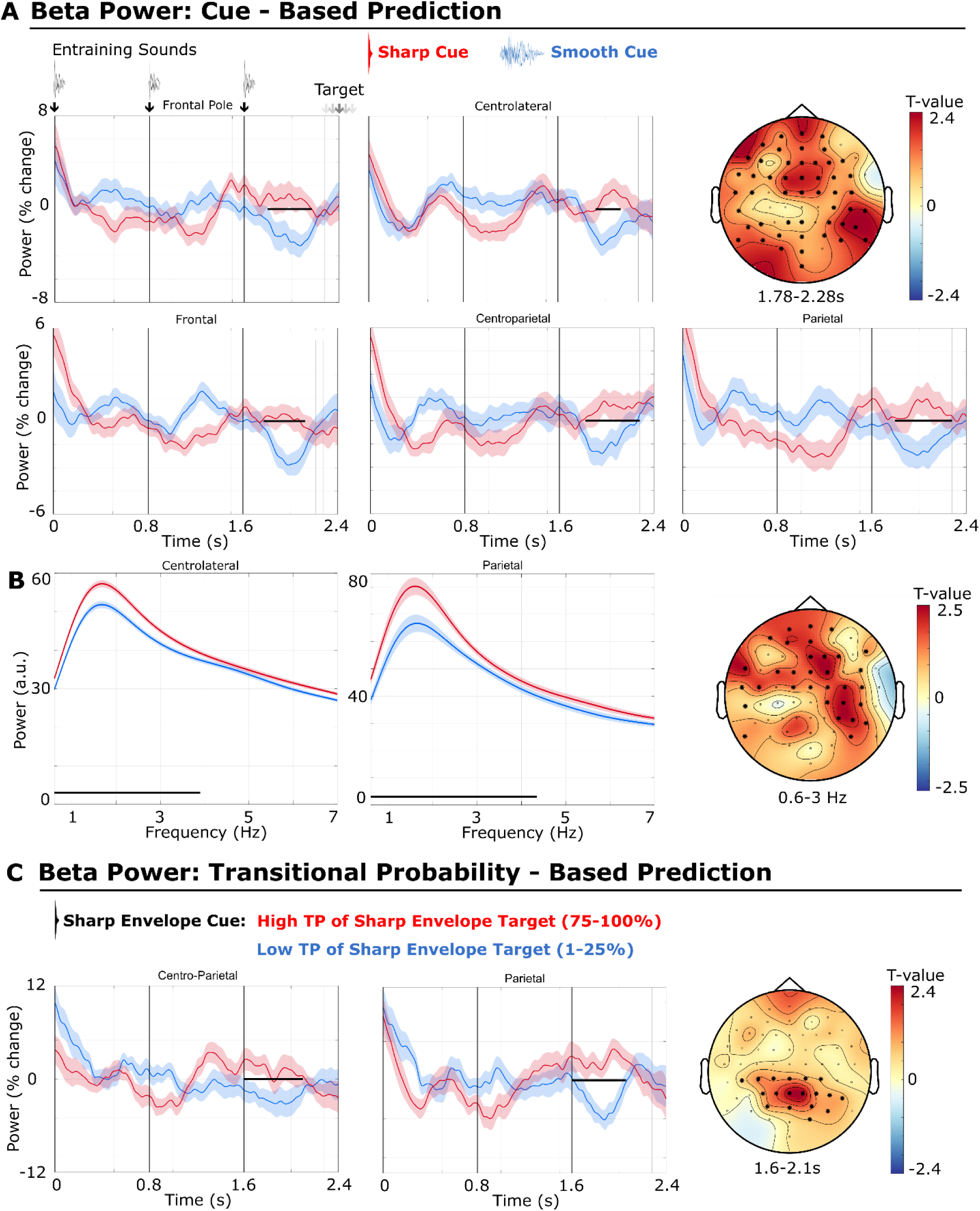
Beta power effects for the sharp versus smooth target prediction based on the cue or the transition probabilities (TP). Shaded areas represent the adjusted standard error (SE) for a within-subject design, according to the Cousineau-Morey correction (O’Brian & Cousineau, 2014). Black lines indicate significant differences in the temporal dimension. The spatial extent of the significant clusters is depicted by bold stars in the topographical plots. The small gray arrow symbols for the target depict multiple possible onsets for the target according to individual P-Centers. (A) Pre-target beta-power time series (percentage change) for sharp (red) versus smooth (blue) cue condition. Pre-target beta power is significantly enhanced for the sharp cue condition. For visualization purposes, beta power was averaged over significant sensors of selected electrode placement sides: Frontal Pole (Fpz, Fp1, Fp2), Frontal (Fz, F1, F2, F4–7), Centrolateral (C4–6, T7), Centroparietal (CP4, CP6, TP7, TP8), Parietal (Pz, P1, P4–8). (B) Spectral modulation of beta-power time series for the sharp (red) versus smooth (blue) cue condition. The spectral modulation of the pre-target beta-power time series is significantly enhanced for the sharp cue condition within the delta (1–3 Hz; topographical plot shown on the right) and theta (4–7 Hz) frequency ranges (data not shown). The spectral modulation of beta power was averaged over significant sensors of selected electrode placement sides: Centrolateral (C4–6, T7), Parietal (P4, P6, P8). (C) Pre-target beta-power time series (percentage change) for high (75th percentile or upper quartile) versus low (25th percentile or lower quartile) transition probability (TP) to receive a sharp target sound. Transition Probabilities reflect the probability of a sharp target sound in a current trial, based on the history of valid versus invalid sharp cue trials (for details, see “Methods”). Differences in TP are shown for the sharp cue condition, to control for perceptual confounds caused by physical differences between the sharp and the smooth sound cue. Beta power is modulated in the same direction, showing increased beta power for a high (75–100%) TP to receive a sharp target sound. For visualization purposes beta power was averaged over significant sensors of selected electrode placement sides: Centroparietal (CPz, CP1–5, TP7), Parietal (Pz, P1 -4, P6).

The power spectra of the beta-power time series showed a peak at approximately 1.6 Hz for both conditions, revealing that the strongest modulation of the beta-power time series was in the delta range (1–3 Hz), entailing the beat rate (1.25 Hz) of the entraining sound (Figure 3B). Beta-power dynamics showed a significantly stronger modulation at the delta frequency range (Figure 3B) for the sharp cue condition (one-sided cluster-corrected permutation test, n = 19, cluster T-sum = 3229.7459, cluster P-value = 0.0247). This modulation was also apparent for five electrodes for the theta (4–7 Hz) frequency range. These frequency modulations involved central, frontal, and parietal sites (Figure 3B). The effect size for the average power across channels and frequency bins was Cohen’s d = 0.4939.

### Beta-band power correlates with performance in the timing-judgment task

We tested neural-behavioral correlations to reveal the behavioral relevance of the identified pre-target beta-power modulation effects (i.e., the enhancement of beta power for the prediction of a sharp sound), inducing high perceptual timing precision. If beta-power modulations reflect underlying neural processes involved in the prediction of envelope sharpness and perceptual timing precision, then we would expect a positive correlation between timing judgment performance and beta-power t-values (sharp versus smooth cue contrasts at the individual level).

As our initial behavioral index, we selected the sensitivity measure (d-prime) that reflects the performance in the timing judgment task in the valid sharp cue condition. We investigated its relation to the individual beta-power t-values (sharp versus smooth cue condition) via Spearman correlations, restricting the analysis to the time window showing a significant group effect for the pre-target beta power (1.78–2.28 s; see results above). The group-level test was performed with non-parametric cluster-based permutation tests. It showed a significant correlation (cluster t-sum = 5174.5376, cluster P-value = 0.0120) between the d-prime (for the valid sharp cue condition) and the beta-power t-contrast, starting at the earliest at 1.905 s and lasting at the most up to 2.28 s (i.e., shortly before target onset), with an extended topography over frontal, temporal, parietal, and occipital sites (see Figure 4). The average effect size across the cluster (channels and time bins) was rho = 0.5311. Except for two electrodes, all showed a significant positive correlation between individual beta power t-values and detection performance (d-prime for the valid sharp cue condition).

As a second behavioral measure, we were interested in the P-center variability, which increases with decreasing precision in the perception of the temporal location of a sound. We hypothesized a negative correlation between P-center variability and the beta-band power t-contrast (sharp versus smooth cue condition). As expected, individual differences in P-center variability were most pronounced for the smooth sound (see the “Behavioral Results” section). Therefore, we calculated non-parametric Spearman correlations between the individual P-center standard deviation for the smooth sound (a measure of individual precision ability) and the individual beta-power t-values for the sharp versus smooth cue condition. Non-parametric cluster-based permutation tests on the group level revealed significant negative correlations (cluster t-sum = -2651.3616, cluster P-value = 0.0410) starting already at 1.865 s and lasting up to 2.28 s, with a topographic distribution encompassing the frontal, central, and posterior parts of the sensor space (Figure 4). The average effect size across the cluster (channels, time, and frequency bins) was Spearman’s rho = -0.5055 (medium association). The most pronounced effects were found at posterior sites, followed by anterior sites.

**Figure 4.**
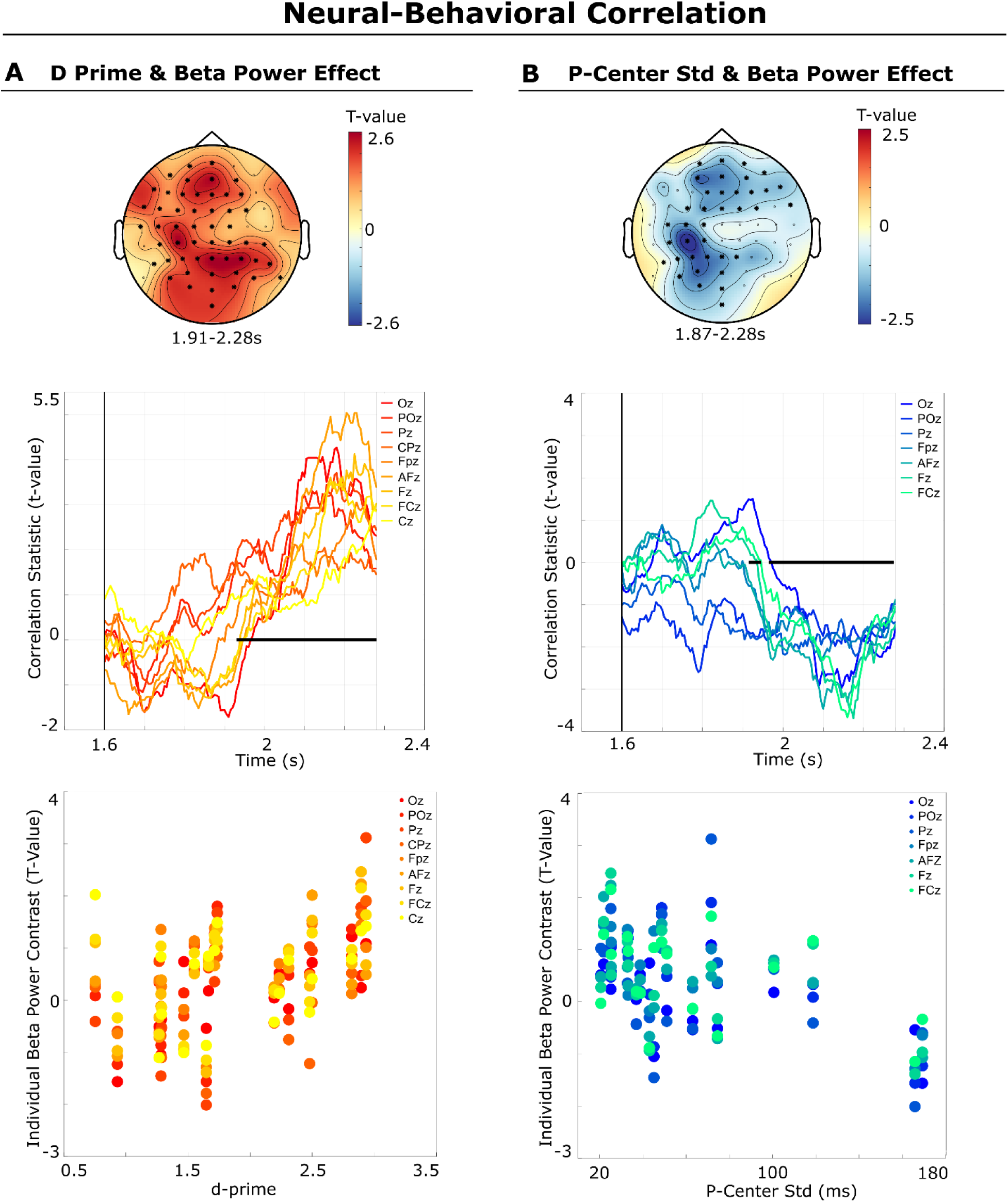
Neural-Behavioral Correlation. Top: topographical distribution of the correlation effects. Middle: correlation time courses of the significant midline sensors. Bottom: correlation at the individual subject level (each dot represents a participant for the significant midline sensors). (A) Correlation between the behavioral d-prime measure for the valid sharp cue condition and the individual beta-power contrast (t-value) between the sharp versus smooth cue conditions. There was a positive correlation between the individual beta-power effect and the d-prime result for the timing judgment task. (B) Negative correlation between the behavioral P-center variability measure (std) for the smooth sound and the individual beta-power contrast (t-values). Together, these results support the hypothesis that pre-target oscillatory beta activity encodes the predicted envelope sharpness and temporal precision of the upcoming target sound.

These results indicate that the participants who showed the strongest beta power modulation by the cue (i.e., larger beta-band power t-value for the sharp cue condition) both performed better on the timing-judgment task (positive correlation) and showed a higher perceptual timing precision when estimating the P-center of the smooth sound (i.e., lower P-center variability; negative correlation).

## Discussion

Music and speech are structured in time by rhythmic modulations in sound intensity. In speech, slow temporal modulations reflect the syllabic (∼5 Hz) and phrasal (0.5 – 2 Hz) rate (Ding et al., 2017, 2016; Goswami and Leong, 2013; Rimmele et al., 2021). In music, slow modulations (0.5–3 Hz) reflect acoustic fluctuations near the beat rate (Ding et al., 2017), underlying the perception of beat, meter, and grouping (Ding et al., 2017; Gordon, 1987; Large and Palmer, 2002; London, 2012). In the current study, we investigated how predictive information about low-level features of acoustic stimuli—namely, the sharpness of the envelope—might serve the prediction of a target that is embedded in such a musical beat. The goal was to reveal neural processes underlying the prediction of the envelope sharpness and how this affects temporal processing. We argue that the acoustic envelope shape is closely related to perceptual timing precision (Danielsen et al., 2022, 2019) and believe that neural beta activity encodes the prediction of envelope sharpness. This neural process might also be involved in top-down control of neural entrainment to quasi-periodic rhythmic events in music and speech.

Flexibility in perceptual timing precision allows us to perceive regularity in acoustic information that does not show a strictly periodic pattern or that has a shape that renders its perceived temporal position indistinct, as can be the case in language (Bonnet et al., 2024; Poeppel and Assaneo, 2020) and music (Danielsen, 2010; Danielsen et al., 2019). Although neural entrainment is often demonstrated for strictly isochronous stimuli, the neural mechanisms seem to tolerate a certain amount of temporal variability (Lakatos et al., 2019). Such tolerance for temporal variability is also more adaptive, given that most environmental events do not follow strict isochrony (Bonnet et al., 2024) and tend to vary in their acoustic envelope shape. Nevertheless, studies of neural tracking to quasi-periodic sounds that consider the predictability of the acoustic envelope sharpness and related effects on temporal perception are lacking. With the current study, we intend to close this gap.

We designed a timing-judgment task in which we utilized sound cues to manipulate expectations about the sharpness of the envelope of an imminent target sound following a short beat sequence. In this way, we induced different predictions regarding the perceptual timing precision of the target, which in turn affected temporal processing during the timing judgment task. We show that beta activity encodes the expected sharpness of the acoustic envelope of upcoming auditory events, and, importantly, that this prediction process is under top-down control.

### The expected sharpness of the acoustic envelope modulates temporal perception

The results show that perceptual sensitivity was affected by the cue, supplying clear evidence that the sharpness of the acoustic envelope indicated by the cue was behaviorally relevant and had an impact on the temporal perception of the target.

We used acoustic features (i.e., attack and duration) to manipulate the envelope sharpness of sounds. Our hypothesis was that we would thereby induce different levels of perceptual timing precision in the processing of the target sound. To ensure that the acoustic features had the intended effect on temporal perception, we measured the P-center location and variability, which reflect how precisely a listener locates a sound in time. We replicated the results of previous studies (Câmara et al., 2020; Danielsen et al., 2019), as the sound with a sharp acoustic envelope had a significantly lower perceptual variability (i.e., a higher perceptual timing precision) than did the smooth sound. These results verify that the target sounds affected perceptual timing precision in the intended direction.

Furthermore, the relevance of the pre-target cue for task performance was a prerequisite for our argument that the observed neural process is under top-down control. The results show that timing judgment performance was indeed modulated by cue validity for target sounds with a sharp envelope. This means that expecting a target with a smooth envelope was detrimental to the judgment of the timing when the actual target was a sharp envelope sound that required high perceptual timing precision. In contrast, cue validity did not significantly affect behavioral performance for target sounds with a smooth envelope, which induce low perceptual timing precision. That is, a cue that erroneously signaled a sharp envelope target was not detrimental to the subsequent temporal processing of a smooth target. The performance enhancement by the valid sharp cue is a pivotal finding because it shows that the top-down prediction of the sharpness of the envelope affects temporal processing. This indicates that the neural prediction of the envelope shape also incorporates a prediction of the perceptual timing precision of the target sound.

Interestingly, the post-experiment self-report (an open-ended questionnaire) contradicted the behavioral and neurophysiological results. Most of the participants (79%) stated that they did not use the cue to solve the timing judgment task. This suggests that most participants were not consciously aware that the perceptual system uses the cue information to predict the acoustic shape of the target and, in turn, the requisite perceptual temporal precision for performing the task. On the other hand, most people (68%) reported that it was easier to judge the timing of the sharp target sound than the smooth target sound. This is interesting because it coincides with the behavioral advantage of the cue, which only arose for the sharp target sound. Altogether, our findings show that the neural system exploited the information conveyed by the cue to predict different levels of envelope sharpness and the related perceptual timing precision of sounds in a top-down manner while tracking a quasi-periodic beat.

### Beta-band power encodes predictions about the acoustic envelope sharpness and temporal precision

At the neural level, the results replicated previous findings that modulations in beta-band power synchronize with a regular beat (Fujioka et al., 2015; Merchant et al., 2015) and encode temporal predictions (Foldal et al., 2023; Morillon and Baillet, 2017). Importantly, this study revealed that beta power is modulated in a top-down manner by the cue, enabling the neural system to flexibly adapt to different acoustic envelope shapes in accordance with task-specific demands, thereby enhancing sensory selection and performance.

The ongoing debate about whether oscillatory entrainment is instrumental to temporal prediction revolves around the central question of whether it can be actively modulated in a top-down fashion. Growing evidence indicates that rhythmic stimuli with a regular structure facilitate temporal processing and enable behavioral benefits thanks to temporal anticipation (Correa and Nobre, 2008; Henry et al., 2014; Henry and Obleser, 2012; Jones, 2018). Neural oscillations reflect rhythmic fluctuations of neural excitability and are, therefore, excellent candidates for a mechanism underlying temporal prediction via the phase alignment of slow oscillations to external events (Henry et al., 2014; Henry and Obleser, 2012; Kösem et al., 2014; Obleser and Kayser, 2019; Suess et al., 2021). This makes oscillatory entrainment an elegant neurophysiological implementation of the dynamic attending theory (Haegens and Zion Golumbic, 2018; Jones, 2018; Lakatos et al., 2019). However, phase synchronization could be caused via bottom-up-driven evoked responses to the regular stimuli that simply reflect the periodicity of the events (Suess et al., 2021; Zoefel and VanRullen, 2016). This makes it difficult to determine whether neural entrainment is functionally relevant to sensory selection (but see Kösem et al., 2014; Zoefel and VanRullen, 2016). Therefore, the key question becomes whether top-down factors can actively control neural entrainment (Haegens and Zion Golumbic, 2018; Suess et al., 2021).

In an effort to synthesize current debates about temporal prediction, Rimmele et al. (2018) argued for a neuronal representation that enables a simultaneous bottom-up (stimulus-driven) and top-down (motor- or language-driven) phase reset of low-frequency oscillations in the auditory cortex that is carried by beta oscillations. Likewise, with reference to the active sensing framework, it has been proposed that a top-down predictive phase reset of low-frequency (e.g., delta) oscillations (i.e., an anticipatory phase reset) could be implemented via the motor system’s encoding of temporal predictions in beta-band activity (Canavier, 2015; Lakatos et al., 2019; Merchant et al., 2015; Rimmele et al., 2018). In line with this idea, Large and Snyder (2009) proposed that beta bursts that synchronize entrain to a rhythm might reflect the information transfer between motor and auditory cortices during rhythm perception. One suggested scenario is a phase reset being implemented by the corollary discharge from the motor command that generates a sensory sampling action, informing the sensory region about the temporal arrival of the sensory input (Lakatos et al., 2019). The phase to which the oscillations are reset/entrained is likely conveying predictions about anticipated features of the external object, like entrainment during selective attention (Lakatos et al., 2013; “spectrotemporal filter”). In the same vein, Cannon and Patel (2021) proposed a causal role for the motor system in beat-based temporal predictions and suggested that the prediction of beat timing is communicated through the dorsal auditory pathway connecting the premotor, parietal, and temporal cortices.

The results of the current study provide support for the active sensing framework, since there is clear evidence that neural predictions related to acoustic envelope sharpness are transmitted via beta activity in a top-down manner and modulate temporal perception. Modulations in beta power were aligned to the entraining sounds and relatively enhanced if the participant expected a target sound with a sharp compared to a smooth envelope, and this occurred up to several hundred milliseconds before target onset. Therefore, the modulation of beta oscillations in our study could represent the predictive process of the motor system that informs sensory regions about the probable acoustic features of future events, which enables a top-down phase reset of low-frequency oscillations in the auditory cortex, as suggested by Rimmele et al. (2018). This process could ensure that neural entrainment phase-adjusts to quasi-periodic stimuli such as speech and music in a predictive manner.

Since we did not perform a cross-frequency coupling analysis, we lack direct evidence for a respective phase reset of delta oscillations caused by beta bursts. But we were able to show that pre-target beta-power dynamics are more strongly modulated in amplitude in the delta (beat) frequency range for higher expected envelope sharpness, suggesting an interaction between beta power and slow-frequency oscillations.

To rule out possible confounds caused by different stimulus properties of the sound cues, we calculated for each trial the likelihood of a sharp target sound based on the transition probabilities of the past trials. This more dynamic measure allowed us to contrast trials with high versus low probability to receive a sharp target sound within the sharp cue condition. Beta power was still modulated in the same direction, with enhanced pre-target beta power for trials with a high versus a low probability to receive a sharp target sound.

This rules out purely bottom-up-driven neural responses for the beta-power results and confirms a top-down contribution to the prediction processes that incorporate the information provided by the cue.

Overall, the results are in line with studies indicating that temporal sensory predictions are implemented via beta oscillations (Foldal et al., 2023; Fujioka et al., 2015; Iversen et al., 2009; Morillon and Baillet, 2017; Rimmele et al., 2018; Saleh et al., 2010). Relatedly, Arnal et al. (2015) reported that pre-target beta power was enhanced for correctly detected targets in a timing judgment task, along with a significantly enhanced cross-frequency coupling between pre-target delta phase and beta power.

In addition, brain–behavior correlations indicated that the participants who showed a higher beta-power modulation for sharp versus smooth sound cues both performed better on the timing judgment task of the sharp sound (i.e., larger d-prime; positive correlation) and were more precise at estimating the P-center of the smooth sound (i.e., lower P-center variability; negative correlation). This provides important support for the behavioral relevance of neural beta power for predicting beat timing and the temporal precision of sounds in a beat sequence.

Moreover, beta-band activity was modulated most strongly in the delta range (1–3 Hz), which entails the frequency of the entraining isochronous sounds (1.25 Hz) preceding the target. Importantly, this spectral modulation was relatively enhanced for the sharp compared to the smooth cue condition within the delta and theta (4–7 Hz) frequency ranges during the last entrainment interval before the target. These results are in line with the co-modulation of delta and beta oscillations carrying temporal predictions found by Morrillon et al. (2017). The authors showed that top-down influences via the motor system sharpen sensory processing by encoding temporal predictions via beta oscillations (Morillon and Baillet, 2017). Beta oscillations were directed toward sensory regions and coupled to delta oscillations. The authors argued for the central role of beta oscillations in representing temporal information and encoding top-down predictions, realized via a covert form of active sensing with motor areas providing contextual information to sensory regions (Morillon and Baillet, 2017). Crucially, in the present study, we demonstrate that beta oscillations also convey information about the predicted envelope sharpness, and therefore also the requisite perceptual timing precision, of a sound in a beat sequence in a top-down manner.

In support of the active sensing model, beat perception has been shown to engage the motor cortico-basal ganglia-thalamo-cortical (mCBGT) circuit, with the supplementary motor area (SMA), dorsal striatum, and putamen as important nodes (Cannon and Patel, 2021). This is complemented by studies examining single-cell and microcircuit activity in macaques (Merchant et al., 2015). The idea of the involvement of the motor system fits nicely with the topography of the beta-power results in the current study, which next to fronto-central areas, encompasses predominantly the central and parietal regions of the scalp. However, since we did not source-localize the effect, the interpretation of the scalp topography remains somewhat speculative. Nevertheless, beta oscillations are prominent in the motor system (Pfurtscheller and Lopes da Silva, 1999), and our findings concur with the theoretical framework of active perceptual inference. In addition, the post-experiment self-report demonstrated that 53% of the participants probably had an urge to move to the beat to solve the task.

A recent study (Persici et al., 2023) revealed that the modulation of EEG beta and gamma (> 30 Hz) activity in response to a rhythm was correlated to grammar abilities in children, supporting the idea that neural beat tracking may be an underlying mechanism that is important for both language and music processing. This is in line with studies showing an impairment in the ability to synchronize movements to a beat in children with dyslexia (Colling et al., 2017), who also have been shown to have difficulties to detect the attack (rise time) of sounds (Goswami and Leong, 2013; Huss et al., 2011). Envelope sharpness has been shown to modulate speech tracking (Doelling et al., 2014; Oganian and Chang, 2019) and the precision of beat synchronization, which has been shown in former studies (Danielsen et al., 2022, 2019) and was replicated with the behavioral data in the current study. Therefore, we believe that the neurophysiological results of our study - the encoding of the expected envelope sharpness in the pre-stimulus beta power - could represent a common predictive mechanism that supports both neural speech and beat tracking (or entrainment) in music.

### Limitations and future research

Fujioka et al. (2012) source-localized beta activity to the auditory cortex and found that the beta-event-related synchronization (ERS, power increase shortly before the sound onset) represented the predicted timing of the next sound in a beat sequence. We did not perform a source localization of the beta activity (i.e., the analysis of the ERS and event-related desynchronization [ERD]) in relation to the entraining sounds (0–1.16 s), as this was beyond the scope of this paper. If the listener expected a sharp target sound, then beta-power time series were aligned to the entraining sounds for a subset of temporal and parietal electrodes, with a power increase before the next entraining sounds and a power decrease after the sound, resembling the ERS and ERD described by Fujioka et al. (2015, 2012). However, for the condition inducing a prediction of a smooth target sound, the alignment of the beta activity was quite different, with a beta-power peak halfway between the first and second, and second and third, entraining sounds (around 0.4 s and 1.2 s) for several anterior sensors. This is a highly interesting observation and should be further investigated in a future study conducting EEG source analysis.

Studies investigating rhythm-based and cue-based temporal expectations have shown functional dissociations in behavior and provided evidence that both processes might rely on distinct brain circuits (Breska and Ivry, 2020; Nobre and van Ede, 2023, 2018). In the current study, we did not intend to dissociate cue-based and rhythm-based temporal predictions. Accordingly, the task design does not allow for an investigation of a possible dissociation between these cognitive mechanisms. Our goal was to modulate the beat-based neural tracking process in a top-down fashion via a cue. Still, multiple distinct mechanisms might have been at play and acted in synergy, creating the behavioral benefit of the cue in a task that involves rhythmic expectations.

The results could also be interpreted in light of the dynamic attending theory (Jones, 2018; Large and Jones, 1999), which suggests that attentional processes are inherently dynamic and align to the temporal patterns of external events. Large and Jones (1999) showed how external rhythms might entrain neural slow oscillations by aligning the phases to optimal time points to direct selective attention. In their computational model, the attentional focus increases with certainty and decreases or widens with more variability (uncertainty) in an external rhythm. Jones suggested that, in addition to phase, the amplitude of induced beta is coupled to the phase of the slow oscillations, representing attending pulses that precede the events (Jones, 2018). Similarly, we found a modulation of beta activity depending on the envelope sharpness of the target sound. However, since our paradigm precludes investigation of the phase of slow entrained oscillations due to the evoked responses at the entrainment rate, we cannot directly test predictions by DAT related to the phase synchronization of delta oscillations and the modulation of neural entrainment in the low frequency range.

Another limitation of the study might be that targets differed not only in envelope sharpness and P-center variability but also in P-center location and spectral features (center frequency). Target sounds were P-center aligned to the entraining sounds. The later the P-center location, the earlier the presentation of the sound in order to P-center align it with the foregoing entraining sounds. The neural differences observed for the two levels of envelope sharpness might thus be confounded by different expectations regarding the sound onset and spectral properties (e.g. higher center frequency for the sharp sound) of the sound. However, pre-target beta power was increased for the sharp cue condition, which generally had later sound onsets than the smooth cue (due to the P-center alignment, the smooth sound had to be presented earlier to compensate for the relatively late P-center location). Therefore, a possible confound due to different onset times of the target seems unlikely; moreover, the behavioral finding indicating detrimental effects for only the target sound with a sharp envelope argues against it. We thus believe that the pre-target modulations in beta activity were caused by different predictions of envelope sharpness and temporal precision of the target sound. Danielsen et al. (2019) systematically investigated the influence of all three acoustic features on the P-center variability and found that only attack and duration had a significant influence on the variability of the P-center (here referred to as perceptual timing precision) and not frequency. We therefore believe that the prediction of the attack and duration, i.e. acoustic features that determine the envelope sharpness of sounds, were contributing to the temporal perception effects we found in the current study. To rule out spectral features as a possible confounding factor, a future study could control for the center frequency of the stimuli and only change the shape of the attack of the sounds in order to manipulate P-center variability.

### Conclusions

Taken together, our findings show that the brain exploits the sharpness of the acoustic envelope to predict different levels of perceptual timing precision of sounds in a quasi-periodic sequence of events, and that this process is at least partly under top-down control. Importantly, the results reveal that beta activity is involved in the neural tracking of not only strictly isochronous events but also quasi-periodic patterns, which predominate in speech and many musical styles (Clarke, 1985; Haugen and Danielsen, 2020; Johansson, 2017; Kvifte, 2007; Polak et al., 2018).

The pre-stimulus modulation in beta oscillations could reflect a predictive mechanism that dynamically adjusts neural entrainment to expected acoustic landmarks (sharp envelope events). In line with the active sensing framework (Lakatos et al., 2019; Morillon et al., 2019; Rimmele et al., 2018) and studies relating beta activity to beat perception and temporal predictions (Arnal et al., 2015; Fujioka et al., 2015; Merchant et al., 2015), this could possibly be realized via beta bursts that cause a phase adjustment of slow oscillations that track the syllable rate in speech or the beat in music.

## Methods and Materials

### Participants

Participants were recruited at the University of Oslo. In total, 28 healthy volunteers participated in the study after giving written informed consent. They were compensated with gift cards worth 300 NOK. Beforehand, the Department of Psychology’s internal research ethics committee had approved the study, which was conducted in agreement with the Declaration of Helsinki. Nine participants were excluded because their performance on the timing judgment task was not significantly higher than chance level (see “Behavioral Analysis” below). The final sample included 19 subjects (13 female, all right-handed, mean age = 25.79 years, std = 3.49 years). All reported normal vision and hearing, no central nervous system damage or disorder, and no cognitive difficulties. They also reported that they did not receive any psychiatric treatment or medication for illness at the time of participation.

### Experimental Task and Stimuli

Using a cueing paradigm, we manipulated the prediction of the acoustic envelope shape (sharp vs. smooth) of the target sound. Both the cue and the target sound were embedded in an isochronous sound sequence (Figure 1). The cue was either a sharp (50%) or smooth (50%) envelope sound (further described below), presented at the beginning of each trial. The sound cue indicated the envelope shape of the target sound. In the case of a valid cue (68% of trials), the target sound was identical to the cue, whereas in the invalid condition, it was not identical (32% of trials).

The target sounds followed three isochronous entraining sounds that were physically constant across all conditions. Participants had to judge the timing of the target sound— whether it was on time or delayed—in a two-alternative forced-choice task. The ISI between all three entraining sounds was always 800 ms, resulting in a beat frequency of 1.25 Hz (Figure 1). The ISIs between the cue or target and the entraining sounds were adapted so as to ensure perceptual isochrony via a P-center alignment of the stimuli (explained in detail further below).

To induce high or low perceptual timing precision, we utilized sounds with two different acoustic envelope shapes that induce either a high or a low P-center variability, respectively (Câmara et al., 2020; Danielsen et al., 2019). We refer to these sounds as sharp or smooth sounds. These differently shaped sounds were originally designed to investigate acoustic factors influencing P-center location and variability (Câmara et al., 2020; Danielsen et al., 2019). P-center locations are reported in ms relative to the physical onset of the sound. The noise sound with the highest P-center variability results (group mean std = 28 ms) was selected as a sound inducing low perceptual timing precision and referred to as a smooth sound. This sound had a gradual attack (50 ms rise time), a long duration (400 ms), and a center frequency of 100 Hz (Danielsen et al., 2019). The sound with the lowest P-center variability (group mean P-center std = 2 ms) was selected as a sound inducing high perceptual timing precision and referred to as a sharp sound. This sound had a sharp attack (2 ms rise time), a short duration (40 ms), and a spectral centroid of 2370 Hz (Câmara et al., 2020). The three entraining sounds following the cue sound had a duration of 100 ms, a rise time of 3 ms, and a center frequency of 100 Hz, and they had been shown to induce a P-center variability of std = 7.78 ms (Danielsen et al., 2019), which is between the sharp and the smooth sounds.

Except for the sharp sound, where we assumed a P-center location of 0 ms based on previous studies (Câmara et al., 2020), all sound types were aligned to participant-specific P-center locations (see the procedure for the P-center estimation below) to ensure that the sound series were perceived as isochronous. For example, for a participant with a P-center location of 15 ms for the entraining sound, the onset of the sound was adjusted to be 15 ms earlier. This particular case would then result in an ISI of 785 ms between, for example, the sharp sound cue (P-center of 0 ms) and the following entraining sound but would supposedly induce a perceived isochrony of 800 ms. Given its P-center location of 0 ms (corresponding to the physical onset of the sound), the sharp target sound occurs at approximately 2.4 s for the on-time condition. The target sound was also P-center aligned relative to the entraining sounds, resulting in possible earlier or later target onsets.

Overall, and for each factor combination, 50% of the target sounds were on time and 50% were delayed, resulting in a 2 x 2 x 2 factorial design, with factors envelope sharpness (sharp/smooth) x cue validity (valid/invalid) x target timing (on time/delayed) and eight conditions in total (Figure 1). The sound cues were presented in a pseudorandom order to prevent repetition in successive trials.

Participants indicated whether the target sound was on time (i.e., isochronous to the preceding entraining sounds) by pressing the left green button of a response box (Cedrus Response Pad, Model RB-740, www.cedrus.com) with the index finger of their right hand, or delayed by pressing the right red button with the middle finger of their right hand. Participants were given 1.3 s to respond before the next trial was automatically initiated. The inter-trial intervals (ITI) were uniformly distributed between 1.2 and 2.2 s, with a mean length of 1.7 s. Participants were asked to fixate on a central black cross on a gray background that remained on the screen during the entire experimental block.

The target delay threshold was determined individually for each participant after one initial training block consisting of 100 trials. The training block started with a delay of 90 ms for the sharp target sound and 100 ms for the smooth target sound. After the training block, the threshold was automatically adapted. A one-sided binomial test was used to determine whether the performance (accuracy) was significantly higher than chance level (significantly higher than 50% correct). The target delay was increased if the performance was too low (by 30 ms if the accuracy was below 30%, by 20 ms if the accuracy was below 50%, or by 10 ms if the accuracy was above 50% but not significantly higher than chance level). If the performance was too high (approaching 100% accuracy), then the delay was reduced (by 20 ms if the accuracy was above 90% or by 10 ms if the accuracy was above 80%) to avoid behavioral ceiling effects. Individually adjusted temporal delays were held constant across trials and blocks for the main experimental task. The main experimental task consisted of three blocks of 100 trials each, resulting in 300 trials.

### Estimation of the Loudness of Stimuli

In a pilot experiment, we asked 10 participants (who did not participate in the main experiment) to judge whether the loudness level of the sharp and smooth sounds was perceptually similar to the entraining sound. Participants were allowed to increase or decrease the loudness level of the sharp or smooth sound until it was perceptually similar to the loudness of the entraining sounds. The mean of the amplitude factors for the sharp and smooth sounds across all 10 participants was used as the new amplitude factor for the main experiment (please see stimuli and scripts in the online repository). The sound loudness was at a comfortable level and was constant across participants and blocks.

### P-center Estimation

Before the main task, individual P-centers were estimated via a click alignment task (see Danielsen et al., 2019, for details of the procedure), which is a computerized task to estimate the P-center location and variation (measured in standard deviation). Previous studies used only four (Danielsen et al., 2019) or three (Câmara et al., 2020; Danielsen et al., 2022) trials for the estimation of the P-center location and variability, since pilot tests showed that this was sufficient to provide a good estimate. For the current study, we increased the number of test trials to get reliable estimates for the entraining and the smooth sound. We used nine trials for the smooth sound since this sound creates the highest perceptual variability in a subject and six trials for the entraining sound. We used only three trials for the sharp sound since we did not intend to use the results from the click alignment task for the P-center alignment of the experimental stimuli. A previous study (Câmara et al., 2020) had already shown that the sharp sound had a grand average P-center location of 0 ms and a standard deviation of 2 ms (mean over three trials). Accordingly, we used the respective P-center location for the stimulus alignment in the current study. Please see Figure 2 and Supplementary Table 1 for the individual P-center locations and standard deviations.

### Post-Experiment Self Report

All participants completed a self-report (see Supplementary Figure 1) after finishing the experimental task. The open-ended questionnaire included six open questions regarding their behavioral strategies and their subjectively perceived use of the cue while performing the task.

### Goldsmith Musical Sophistication Index

We collected data for the Goldsmith Musical Sophistication Index (Gold-SMI) questionnaire (Müllensiefen et al., 2014), which was not used for the current paper (see Supplementary Material).

### Procedure

The session started with participants reading and signing the consent form before we set up the EEG recording and the experimental tasks. Auditory stimuli were played at a comfortable volume from two Genelec speakers (model 8030 W) flanking both sides of the screen.

After performing the click-alignment task to estimate the individual P-center location and variability (approximately 5 minutes), participants did a training block (lasting approximately 10 minutes) to estimate the target delay threshold. After a short break, participants performed the three experimental blocks of the main experiment (approximately 12 minutes each). Participants decided if they would like to take short breaks between blocks. The entire session, including EEG preparation, experimental tasks, breaks, and administration of questionnaires, lasted two to three hours.

### EEG Recording

EEG and electro-oculography (EOG) data were recorded using a BioSemi Active Two 64 Ag-AgCl electrode system (BioSemi, Amsterdam, Netherlands). A BioSemi headcap was used to position the 64 electrodes on the scalp of the head, according to the international 10–20 system for electrode placement. Data were sampled at 1024 Hz during online recording. Four external electrodes measured vertical and horizontal eye movements (EOG) and were positioned above and below the right eye, and lateral to the participant’s left and right eyes. Two additional electrodes were positioned on the left and right earlobes for later EEG offline re-referencing.

### Behavioral Analysis

We tested whether the performance in the timing-judgment task was significantly higher than chance level (more than 50% correct detections) using a one-sided binomial test for the accuracy of the valid sharp cue condition and the valid smooth cue condition. If the participant failed to perform significantly better than chance level (p>0.05) in one of the conditions, they were excluded from the analysis. Due to this criterion, nine subjects performing significantly below chance level were excluded, resulting in the final sample of 19 participants.

To quantify the behavioral performance of the timing-judgment task, we calculated the d-prime for each condition based on correct and incorrect responses, which is a performance measure for perceptual sensitivity based on signal detection theory (Stanislaw and Todorov, 1999). We applied a log-linear rule, adding 0.5 to both the number of hits and false alarms and 1 to the number of signal (sum of hits and misses) and noise (sum of correct rejections and false alarms) trials (Stanislaw and Todorov, 1999), before calculating the adjusted Hit and false alarm rates (FA).

By taking the inverse of the standard normal cumulative distribution function 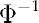 (z-score) for the Hit and FA rates, the d-prime (d’) was calculated as follows:

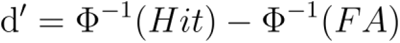

This framework also allows for estimating the response bias separately from sensitivity, which was calculated as follows:

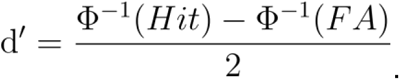

The d-prime and response bias was calculated for all factor combinations: sharp-valid cue, sharp-invalid cue, smooth-valid cue, and smooth-invalid cue, resulting in a 2 x 2 (envelope sharpness x cue validity) factorial design. Target reaction times were analyzed only for the correct responses for all four conditions.

### EEG analysis

#### Data Preprocessing

The focus of the analysis of the neurophysiological data was to investigate whether the factor envelope sharpness (sharp versus smooth cue) affected the neural processing of the Entraining sounds and pre-target oscillatory activity. This modulation might mirror predictive processes optimizing temporal perception. Therefore, we concentrated on the comparison of the sharp versus smooth cue condition, pooling over the two factors target timing and cue validity (both are unknown to the participant before the target arrives), including only trials with correct responses to reduce noise (e.g., attention not on the task, etc.).

EEG data were analyzed offline using custom-written scripts and the Fieldtrip toolbox (Oostenveld et al., 2011) in MATLAB (R2020a, Mathworks Inc., Natick, MA, USA). Continuous EEG data were high-pass filtered with a zero-phase Butterworth IIR filter with a half-amplitude cut-off at 0.1 Hz (18dB/oct roll-off) to remove slow drifts in the data and down-sampled to 1000 Hz. The down-sampling was performed via the Fieldtrip toolbox, which in turn makes use of the MATLAB function “decimate,” incorporating a 400 Hz low-pass Chebyshev Type I impulse response (IIR) filter of order 8 that is applied to the data before down-sampling to prevent anti-aliasing. The data were offline referenced to linked earlobes. Noisy segments and bad channels in the continuous EEG data were defined by visual inspection (i.e., muscle artifacts). Following removal of bad channels, an independent component analysis (ICA) was used to identify and remove blinks and horizontal eye movements in the continuous EEG data. Removed ocular components were identified by visual inspection.

The data were referenced to a common average, segmented into epochs of -0.8 to 3.2 s, and time-locked relative to the onset of the first entraining sound. Epochs containing artifacts were removed. Since the different stimuli types were P-center aligned, the alignment to the first entraining sound ensured that trials could be compared across participants. Note that the three entraining sounds, following the cue and preceding the target sound, were presented at a constant rate of 1.25 Hz (SOA 0.8 s) and were physically constant across all conditions. Epoched data were demeaned and detrended.

#### Frequency Analysis

Time-frequency resolved power was calculated via wavelet transform (Morlet wavelets of 3 cycles) for the entire epoch (-800 to 3200 ms) and the beta frequency range (15–25 Hz) in steps of 1 Hz with a sliding window in steps of 5 ms. Single-trial power values were baseline corrected relative to the power mean across the entire epoch (0–2.4 s) for each frequency bin and expressed in percentage of change (Fujioka et al., 2015, 2012; Fujioka and Ross, 2017). Power values were averaged across frequencies of the beta band (15–25 Hz) to retain beta-power time series (Fujioka and Ross, 2017). Beta-power time series were smoothed with a moving average of 50 samples (equivalent to 250 ms, using the MATLAB function “smoothdata”). The average beta-power time series across trials per participant was calculated as input for the statistical analysis. The time window of interest was the interval between the last (third) entraining sound and the target onset, lasting from 1.6 s to approximately 2.4 s.

The onset of the target sound varied across conditions and participants. Depending on the individual P-center locations of the LP and the entraining sound, the earliest possible target onset was at 2.28 s. (Please see Supplementary Table 1 for individual P-center locations and standard deviations.) We therefore restricted the statistical group analysis to this time window (1.6–2.28 s) to avoid contamination of activity in the pre-target time period via the evoked response of target sounds in some participants.

To reveal slow spectral modulations of beta-power dynamics, single-trial beta-power time series from the time window of interest (last 800 ms before the first possible target: 1.48– 2.28 ms) were submitted to a Fourier transform using a Hanning window to calculate the power spectrum for 1–7 Hz (delta and theta band), which was afterward averaged across trials.

In order to identify spectral peaks in the power spectrum indicative of oscillatory activity, the power spectrum of the last 800 ms of the entrainment period before target onset was calculated by applying a Fourier transform to the EEG raw data of the respective time window (1.48–2.28 s) using a Hanning window and subsequently averaging power spectra (1–30 Hz) across trials, task conditions, and participants.

### Modeling transitional probabilities

The rationale behind the calculation of transitional probabilities (TP) is to express the likelihood of observing a sharp target sound for the current trial based on the entire history of valid and invalid cue-target transitions.

For the estimation of the TP of a sharp cue trial, we divided the number of observed valid cues by the number of valid and invalid cues, to calculate the probability that the current trial is a valid cue-target transition based on the history of trials (observed cue-target transitions). For a sharp cue trial TP, solely observed sharp cue trials (valid and invalid) were considered and for the estimation of the TP of a smooth cue trial, solely observed smooth cue trials were considered. Hence the TP of a sharp cue trial, expresses the probability of receiving a sharp target sound for the current trial. The complementary TP (invalid divided by valid and invalid) was calculated for smooth cue trials. Accordingly, TP always expresses the probability of receiving a sharp target sound and, in case of smooth cue trials, the probability of an invalid smooth cue to sharp target transition.

Transitional probabilities were modeled with a weighted N-back temporal window for the entire history of trials (1,2,…,k) up to the current trial k (Maheu et al., 2019).

An exponential decay function was applied to downweight older transitions over newer ones, with a half-life of 4.16 trials (observations), computed as follows:

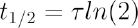

where 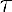 is the exponential time constant: 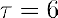, giving larger weight to the most recent trials (Maheu et al., 2019).

This corresponds to a more local integration of observations (leaky integrator). Leaky integrators have been shown to be more biologically plausible in the modeling of brain functions, as opposed to an integration over long time scales with perfect memory (Glaze et al., 2015; Maheu et al., 2019). A short-term integration (half life of 4) has been argued to represent a more deliberate means of searching for local patterns (Maheu et al., 2019). Specifically, the TP of a cue-target transition at a trial k was computed as follows:

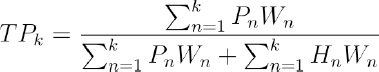

where W, P, and H are vectors of length k (past trials). W is a weighting vector, with elements:

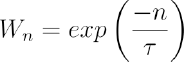

for the k-th past stimulus.

If the current trial entails a sharp cue, then:

1. P elements are defined as 1 for valid sharp cue-target trials (sharp cue to sharp target transitions) and 0 for all other instances (invalid sharp cue to smooth target transitions, valid and invalid smooth cue transitions), and
2. H elements are defined as 1 for sharp invalid cue-target trials (sharp cue to smooth target transition) and 0 for all other instances (valid sharp cue to sharp target, smooth valid and invalid transitions), for the past trials until k.

If the current trial (k) entails a smooth cue, then:

1. P elements are defined as 1 for invalid smooth cue-target trials (smooth cue to sharp target transitions) and 0 for all other instances (valid smooth cue to smooth target transitions, valid and invalid sharp cue transitions), and
2. H elements are defined as 1 for valid smooth cue-target trials (smooth cue to smooth target transitions) and 0 for all other instances (invalid smooth cue to sharp target, sharp valid and invalid transitions), for the past trials until k.

Please note that the weighting vector is applied irrespective of type of cue, corresponding to a temporal weighting.

### Statistical analysis

For the behavioral measures (d-prime and response bias), we conducted two-way repeated-measures ANOVAs on the factors envelope sharpness and cue validity (2 x 2 factorial design). Assumptions of normality were checked via Lilliefors goodness-of-fit test of composite normality and via visually inspecting Q-Q plots. If distributions deviated substantially from normality, non-parametric statistical tests were performed. The sphericity assumption was assessed via Mauchly’s test. If the assumption of sphericity was not met, the Greenhouse-Geisser correction was applied if epsilon<0.75 and the Huynh-Feldt correction was applied if epsilon>0.75. Dependent-samples t-tests were used for the simple contrasts for all conditions of the 2 x 2 factorial design. Simple contrasts for reaction times were also computed for the conditions valid versus invalid (pooled over the levels of the factors envelope sharpness and target timing). P-center locations and standard deviations were compared via non-parametric signed-rank tests for all four conditions of the 2 x 2 (envelope sharpness and cue validity) factorial design (pooled over the condition levels of target timing), since assumptions of normality were violated.

Statistical group analysis of beta-power time series and spectral modulation of beta-power time series for the two conditions of interest (sharp versus smooth cue conditions) were conducted via one-tailed non-parametric cluster-based permutation dependent-samples t-tests, correcting for multiple comparisons (Maris and Oostenveld, 2007), applying 10,000 permutations. Based on previous studies showing that beta power correlates positively with temporal accuracy and prediction (Arnal et al., 2015; Fujioka et al., 2015, 2012; Merchant et al., 2015; Morillon and Baillet, 2017), we had a clear hypothesis about the directionality of the effect: a beta power increase accompanies predicted high envelope sharpness and perceptual timing precision.

Single-trial beta-band power time series were submitted to two-sided independent-samples t-tests contrasting the sharp versus smooth cue conditions to retain individual t-values for the group correlation between neural and behavioral data. Correlations of individual beta-power t-values (sharp versus smooth) with behavioral measures (d-prime and P-center variability) were investigated by performing one-tailed non-parametric

Spearman’s rank correlations using non-parametric cluster-based permutation tests to correct for multiple comparisons, performing 10,000 permutations. Individual d-prime values for the valid sharp envelope cue condition and individual P-center variability results (std) for the smooth sound from the click alignment task were used for this correlation analysis. The correlation analysis was restricted to the pre-target time window of 1.8–2.28 s, according to the results from the statistical group beta-power contrast, showing significant modulation for the factor envelope sharpness (see the “Results” section concerning beta-band power).

To estimate the effect size for cluster-based permutation statistics, we computed average (power or phase) values for each participant and condition across the cluster extent (channels, time bins, and frequency bins, if applicable) and calculated Cohen’s d effect size for within-subjects designs (Lakens, 2013). For correlational analysis, the average Spearman’s rho across the cluster (channels and time bins) is reported as effect size.

## Acknowledgments

We thank Sandra Solli, Olgerta Asko, and Justin London for fruitful discussions, and we thank the volunteers for participating in this study.

## Data sharing plans

Data, documentation, and code used for the analysis will be shared in an online public repository.

