## Supplementary Material for "Predicting the Beat Bin: Beta Oscillations Predict the Envelope Sharpness in a Rhythmic Sequence"

### Predicting the Beat Bin – Beta Oscillations Support Top-Down Prediction of the Temporal Precision of a Rhythmic Event

### Supporting Information Text

#### Materials and Methods

**Goldsmith Musical Sophistication Index**. We included the Goldsmith Musical Sophistication Index (Gold-SMI) questionnaire (Müllensiefen et al., 2014) which is a self-report inventory indicating the level of musical experience. This score incorporates aspects from five sub-scales measuring self-reported active musical engagement, perceptual abilities, musical training, singing abilities, and sophisticated emotional engagement with music. For the current study, we did not use the Gold-SMI results, but the data are available from the authors upon request.

#### Results

**Results of the Post-Experiment Questionnaire**. Participants filled out an open questionnaire to self-report on how they solved the task. 57% of the participants reported that they used a strategy to solve the task and 55% of those used counting as a strategy (31% of the total number of participants). Of all participants, 36% reported to have changed strategies during the experiment, 47% did not feel like guessing, 32% occasionally felt like guessing and 21% felt like guessing with respect to the delay detection task. See the Main Manuscript for the other Results of the self-report.

Date:
Subject ID:

*Post-Experiment Questionnaire*

1. Did you apply a particular strategy to solve the task?

2. Did you change strategies between blocks?

3. Did you move with the beat?

4. Did you consciously use the cue to solve the task? If yes, how did you use it?

5. Did you feel like you were guessing?

6. For which target stimulus was the time judgement easier? (Click or Long sound)

**Fig. S1.** Post-experiment self-report, filled out by the participants.

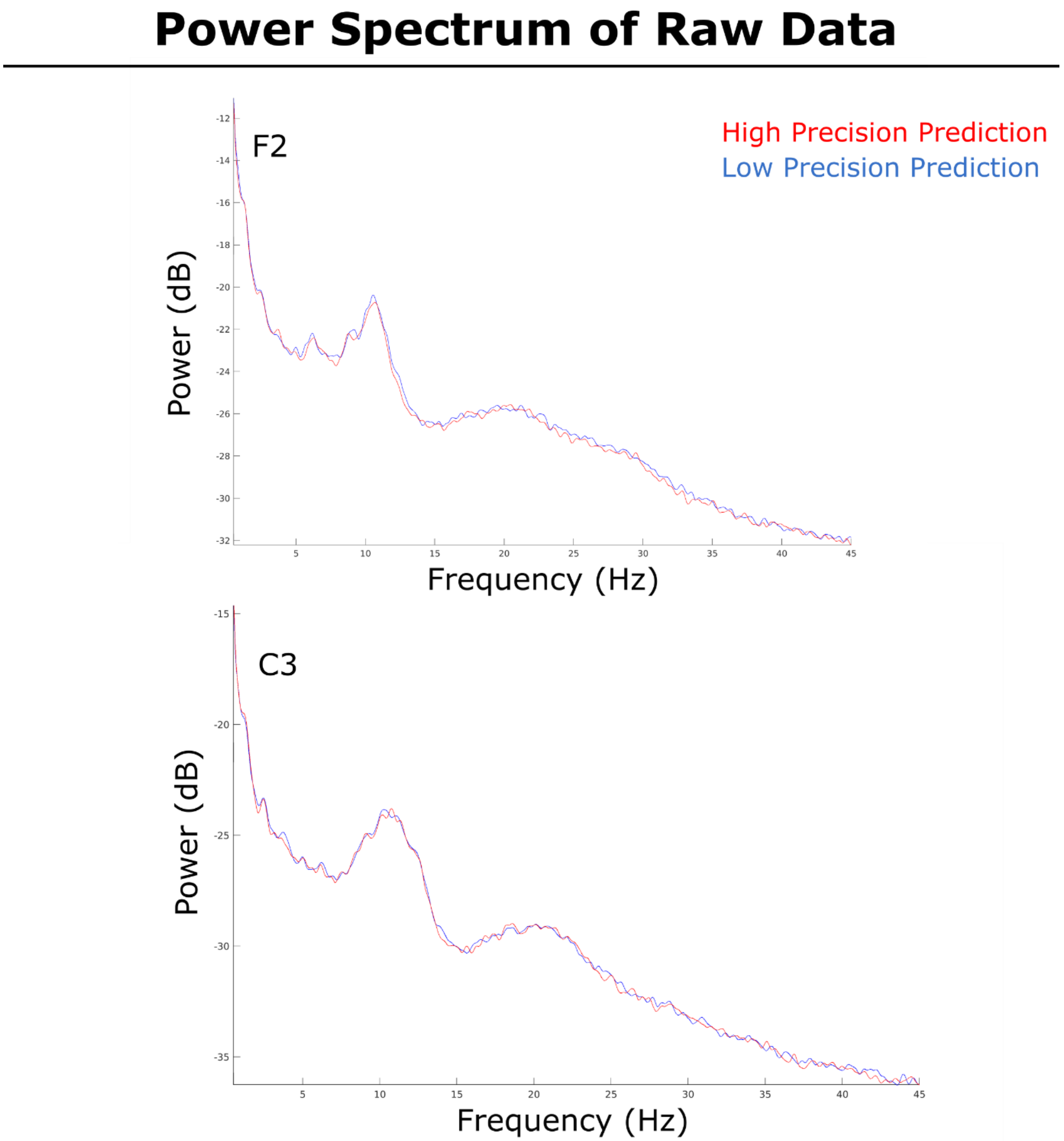


**Fig. S2.** Grand average power spectra for the entire entrainment period (0-2.4 s) for the High- and Low-Precision Prediction condition for frontal (F2) and central (C3) example sensors. The spectra reveal narrowband peaks of power at the theta (4-6 Hz), alpha (8-12 Hz), and beta (15-25 Hz) frequency ranges, indicating the involvement of oscillatory generators.

| Participant | WDB location | FSL location | SLL location | WDB std | FSL std | SLL std |
|---|---|---|---|---|---|---|
| 1 | 17.00 | 26.67 | 57.89 | 34.77 | 37.36 | 38.74 |
| 2 | -29.33 | -19.83 | 6.56 | 31.47 | 29.96 | 74.49 |
| 3 | 9.33 | 2.50 | 56.44 | 38.66 | 60.65 | 33.13 |
| 4 | -13.33 | 9.33 | 44.44 | 19.50 | 53.71 | 51.02 |
| 5 | 6.33 | 60.33 | 94.22 | 15.50 | 47.40 | 71.63 |
| 6 | 17.67 | 27.17 | 63.67 | 12.86 | 27.52 | 32.82 |
| 7 | -71.00 | 14.17 | 19.11 | 89.15 | 47.33 | 169.22 |
| 8 | -2.33 | 15.67 | 56.00 | 13.32 | 22.07 | 43.04 |
| 9 | 12.00 | -8.67 | -3.56 | 9.85 | 38.04 | 165.79 |
| 10 | 37.67 | 29.67 | 45.78 | 40.53 | 27.27 | 45.10 |
| 11 | 22.00 | 8.00 | 44.67 | 6.56 | 24.68 | 25.17 |
| 12 | 18.33 | 19.17 | 68.67 | 30.01 | 48.99 | 100.57 |
| 13 | -20.00 | 18.17 | 47.67 | 30.05 | 23.27 | 36.96 |
| 14 | 8.00 | 60.00 | 65.67 | 1.00 | 109.83 | 63.01 |
| 15 | 12.67 | 1.33 | 35.22 | 4.93 | 27.03 | 21.87 |
| 16 | 56.00 | 74.50 | 81.78 | 33.78 | 21.45 | 48.68 |
| 17 | 9.00 | 2.33 | 40.67 | 3.00 | 10.39 | 25.31 |
| 18 | 6.33 | 15.67 | 31.00 | 3.51 | 19.71 | 20.52 |
| 19 | 33.67 | -8.83 | 112.89 | 141.75 | 135.89 | 118.68 |

**Table S1.** Individual P-Center locations and variability (standard deviation) for all three sound types used in the study.
